# The effective number of shared dimensions: A simple method for revealing shared structure between datasets

**DOI:** 10.1101/2023.07.27.550815

**Authors:** Hamza Giaffar, Camille Rullán Buxó, Mikio Aoi

## Abstract

A number of recent studies have sought to understand the behavior of artificial and biological neural networks by comparing representations across layers, networks and brain areas. Simultaneously, there has been growing interest in using dimensionality of a dataset as a proxy for computational complexity. At the intersection of these topics, studies exploring the dimensionality of shared computational and representational subspaces have relied on model-based methods, but a standard, model-free measure is lacking. Here we present a candidate measure for shared dimensionality that we call the effective number of shared dimensions (ENSD). The ENSD can be applied to data matrices sharing at least one dimension, reduces to the well-known participation ratio when both data sets are equivalent and has a number of other robust and intuitive mathematical properties. Notably, the ENSD can be written as a similarity metric that is a re-scaled version of centered kernel alignment (CKA) but additionally describes the dimensionality of the aligned subspaces. Unlike methods like canonical correlation analysis (CCA), the ENSD is robust to cases where data is sparse or low rank. We demonstrate its utility and computational efficiency by a direct comparison of CKA and ENSD on across-layer similarities in convolutional neural networks as well as by recovering results from recent studies in neuroscience on communication subspaces between brain regions. Finally, we demonstrate how the ENSD and its constituent statistics allow us to perform a variety of multi-modal analyses of multivariate datasets. Specifically, we use connectomic data to probe the alignment of parallel pathways in the fly olfactory system, revealing novel results in the interaction between innate and learned olfactory representations. Altogether, we show that the ENSD is an interpretable and computationally efficient model-free measure of shared dimensionality and that it can be used to probe shared structure in a wide variety of data types.

## 1 Introduction

Modern scientific experiments are generating increasingly high dimensional datasets across multiple modalities. In neuroscience, for example, we can now combine information from electrophysiological recordings from different brain regions (eg. [34, 8]), from the same neurons under different conditions, or neurophysiological recordings with behavioural or environmental data. To unravel the relationships contained therein and build theories from these multi-area and/or multimodal data streams, we might seek to decompose the variability in paired datasets into independent and shared components. The variability shared between neural populations or modalities of data is of particular interest as it can often give useful insight into the computational processes at play. In this work, we develop a method for characterizing shared variability between paired datasets and demonstrate its use in a range of data contexts. While we focus our attention on the neuroscience and machine learning applications in the present work, interest in multi-model data analysis reaches well beyond neuroscience as integration of data from different sources becomes routine in applications across the sciences.

Measuring the size (dimensionality) and describing the organization (geometry) of data are key ways of characterizing multivariate relationships. The geometry of shared subspaces has been studied in a wide range of contexts in neuroscience [32, 41, 24, 40, 13], where its frequently referred to as Representational Similarity Analysis (RSA), and machine learning [20, 37, 25, 36], as well as in the intersection of the two domains [19, 38]. In contrast, the dimensionality of shared subspaces has received comparatively little attention, however a number of recent studies have explored the relationship between the dimensionality of neural responses and that of connected tasks or stimuli, finding that the two are related in theory [15] and in experiment [33]. Model based approaches to measuring shared variability such as those used in the latter study may not generalize easily to other contexts and often require sequential hypothesis testing for each candidate shared dimension. Until now, a simple, model free measure has been lacking.

In this work, we present a candidate model-free measure of the embedding dimensionality of a subspace shared between two datasets, which we call the *effective number of shared dimensions* (ENSD). We show that this measure can also be decomposed to describe geometric features of the shared subspace. The ENSD is a generalization of the participation ratio (PR), a measure of the embedding dimensionality for a single dataset, to two datasets. Since its introduction in atomic spectroscopy, the PR has found use in disciplines from quantum and condensed matter physics to economics, sociology and machine learning. The PR has also found wide use in neuroscience [1, 23, 29, 28, 22], where it has been argued that the PR is a reasonable measure of neural data dimensionality [15]. In one recent theoretical study, the authors formally tied dimensionality (PR) to computational task performance in a model of the fruit fly mushroom body, finding that high dimensional representations facilitate lower classification error for simulated odors [22]. Another study on recurrent spiking networks found that localized connectivity features can strongly influence a network*’*s global dimensionality [29], suggesting that dimensionality is sensitive to aspects of local connectivity.

In this paper, we first explore the properties of the ENSD, including an associated distance metric and measure of eigenvector alignment, demonstrating its use in synthetic data. We then contrast the ENSD with two frequently used techniques for analysing shared structure in paired datasets, finding that the ENSD is especially well suited to sparse data. Finally, we demonstrate how these tools can be used to probe both artificial and biological neural systems via analysis of available experimental data in different modalities. We focus our attention on sparse connectomic data from the fruit fly, where application of the related PR has already yielded insights into neural computations. We conclude with some discussion about areas of future development.

## 2 Background

### The participation ratio

The participation ratio (PR) is a measure of the dimensionality of a space defined by a data matrix. For example, for the *n* × *p* matrix **X**, *n* > *p* we assume that the covariance of the rows of **X** are given by 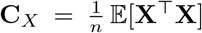. We then denote the eigenvalues of **C**_*X*_ as *λ*_1_ > *λ*_2_ > *…* > *λ*_*p*_. The PR is given by

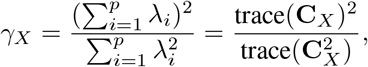

where we can estimate *γ*_*X*_ by noting that 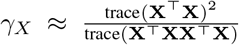. This measure quantifies the dispersion (reciprocal of concentration) of the eigenvalue distribution, giving a more intuitive sense for the effective dimensionality of a data set than many of the existing alternatives [15, 27, 28]. To gain some intuition, consider the set of *p* non-negative eigenvalues *λ*_1_,. .., *λ*_*p*_. By normalizing these values we can write the distribution of the variance over the eigenvectors: 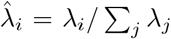. A common measure of the concentration of a distribution is 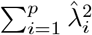, which is constrained to the interval 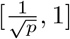. Therefore, 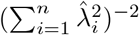 varies on [*1, p*], where 1 is the most concentrated scenario and *p* is the scenario where all eigenvalues are equally distributed and equal to 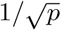. Therefore, if *λ*_i_ are the eigenvalues of **C**_*X*_ then 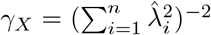 is a measure of how evenly distributed the variance is among all eigenvectors. For several typical eigenspectra, the PR explains between 80-90% of the variance in a dataset [15].

### The effective number of shared dimensions

One may also describe the PR as a matrix inner product. We show this by defining the scaled covariance matrix, 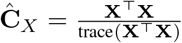. Multiplying this quantity by the participation ratio of **X** we obtain the quantity **G**_*X*_ = *γ*_*X*_ **Ĉ** _*X*_. Then the PR can be expressed as

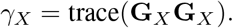

While the PR is a measure of the dimensionality of one dataset, the above representation for the PR suggests a simple generalization that can be applied to the shared dimensionality of two datasets, **X** and **Y**. The only requirement is a shared first dimension, *n*, so we define **Y** to be *n* × *q*. We can now write the expression

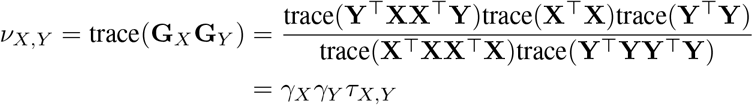

where **G**_*Y*_ is defined as above, *γ*_*X*_ is the participation ratio of **X**, and *γ*_*X,Y*_ = trace(**Ĉ** _*X*_ **Ĉ** _*Y*_).

We call *ν*_*X,Y*_ the **effective number of shared dimensions** (ENSD). Just as the PR is a description of the dispersion of energy across eigenvectors, the ENSD is both a description of how much of the variability in one dataset is explained by the variability in the other and how dispersed that explanation is (Fig.1A). In the following sections, we explore the various properties of this shared dimensionality measure analytically and through a series of illustrative toy examples.

**Figure 1:**
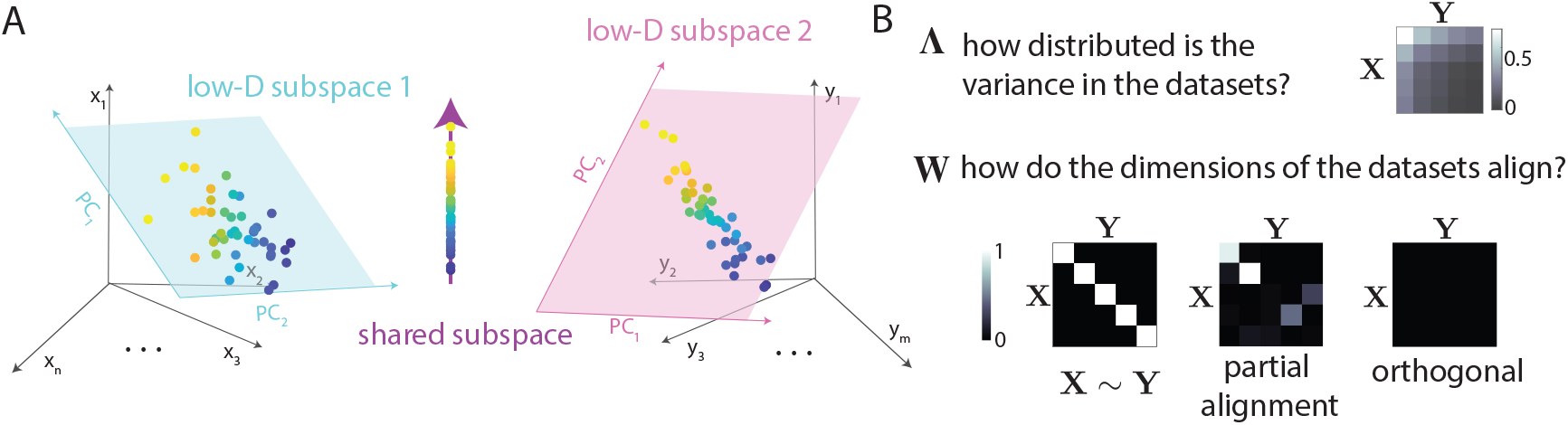
**A**. Schematic illustrating ENSD **B**. Constituent parts of the similarity metric, τ : Λ, a rank-1 matrix of eigenvalue products (top), **W** a matrix of eigenvector overlap (bottom). Examples of **W** are shown for datasets that are i) identical, ii) partially aligned and iii) orthogonal.

### Properties of ENSD (*ν*)

The following are properties of the ENSD which can be proven analytically. The full derivations can be found in the supplementary information (SI).

### Equality with *γ*_*X*_

The ENSD reduces to the PR when both matrices are equivalent up to orthonormal transformation. That is, *ν* (**X, X**) = *γ*_*X*_.

### Upper bound

Just as the PR is upper bounded by min{*n, p*}, the ENSD also admits an upper bound: 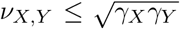. The upper bound is achieved when **X** = **YU**, where **U** is an arbitrary orthonormal matrix, although there may be other conditions under which the upper bound is achieved.

### Corresponding distance metric

Williams et al. [37] noted that inner products can be converted to distances via an arccosine transformation. We can apply such a transformation such that the distance metric

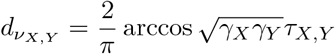

satisfies the equivalence and symmetry requirements as well as the triangle inequality and is therefore a proper distance metric. That we can define such a dissimilarity metric is interesting, first because we can demonstrate a formal relationship between the dimensionality of shared subspaces and a distance in a space defined by a set of equivalence relations. Second, having such an intuitive distance allows this intuition to be put to work in down-stream analyses such as k-nearest neighbors, hierarchical clustering, and a suite of other methods for representational similarity analysis.

We also note that the ENSD allows us to draw a relationship between shared dimensionality and the recently reported Centered Kernel Alignment (CKA) score - a measure that has been applied to measure similarity between e.g. layer activations within or between neural networks [20]. Specifically, the CKA is simply the ENSD normalized by its upper bound: 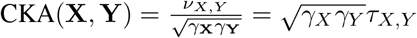. Note that while the CKA can be derived from the ENSD, the inverse is not true. This relationship is explored in more detail below and in the SI.

### Decomposition in terms of eigenvalues and eigenvectors

The ENSD may be rewritten in terms of the eigenvalues and eigenvectors of its constituent matrices. The scaled covariance matrix can be diagonalized as 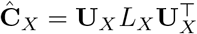, where *L*_*X*_ is the diagonal matrix of normalized eigenvalues 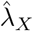 and **U** are the principal axes of the data. We may then write

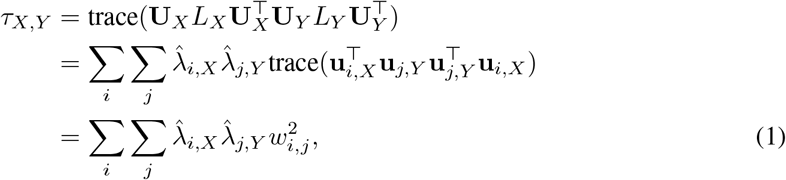

where **u**_*i,X*_ is the *i*^*th*^ eigenvector of **Ĉ** _*X*_, and 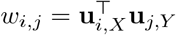. The full derivation is included in the SI. This expression explicitly shows how *ν*_*X*,Y_ is a function of both the eigenvalues of **Ĉ** _*X*_ and **Ĉ** _*Y*_ and the inner products of their constituent eigenvectors. For **X** = **Y**, the entries of 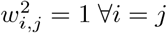 and 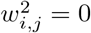 otherwise, and we therefore find 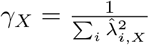.

Since every term in (1) is indexed by *i* and *j* we can rewrite them as entries in the *i, j*^*th*^ position in a matrix. Let 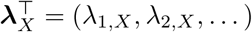, and

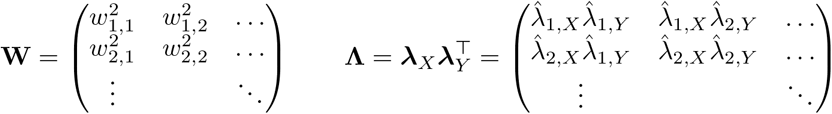

then 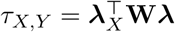 and therefore the ENSD can be expressed exclusively in terms of eigenvalues of eigenvectors:

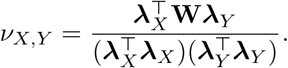

This way of expressing *τ*_*X,Y*_ allows for an intuitive interpretation of this factor: it captures the overlap between the two subspaces via **W**, which is then scaled by the similarities of the eigenspectra, **Λ** (Fig.1B). This similarity measure is then weighted by the individual dimensionalities of the two subspaces in the equation for *ν*_*X,Y*_ to arrive at a final numerical estimate for the dimensionality of the shared subspace.

## 3 Probing ENSD using synthetic data

Here we present a series of toy problems that have been selected to facilitate the development of an intuition for the behavior and properties of ENSD and its relation to its constituent factors, 1 and *τ*. The purpose of this section is to assist in the interpretation of data analysis results using ENSD.

### Shared basis vectors

As a simple example, we construct a scenario where we define the number of shared dimensions between **X** and **Y** to be an integer *r*. Suppose the singular value decompositions of these matrices are given by the *n* × *p* matrix 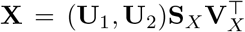, where **S**_*X*_ = diag(*σ*_*X*,1_,. .., *σ*_*X,p*_) and the *n*×*q* matrix 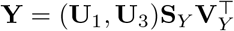, where **S**_*Y*_ = diag(*σ*_Y,1_,. .., *σ*_*Y,q*_), where the key feature we impose is that **U**_i_ ⊥ **U**_j_ for *i* ≠ *j*. The shared subspace is defined by the *n* × *r* matrix **U**_1_, while **U**_2_ is *n* × (*p − r*) and **U**_3_ is *n* × (*q − r*). The *ν*_*X,Y*_ can be written

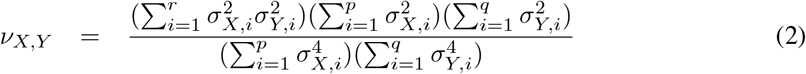

We see from (2) that the shared dimensionality depends on both the exact number of shared eigenvectors *r* and the concentration of the spectra (i.e. the decay of the singular values) of **X** and **Y**. In the case where the dimensionality of **X** and **Y** are maximized (i.e. *γ*_*X*_ = *p, γ*_*Y*_ = *q*), all singular values are equal. If we define 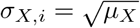 and 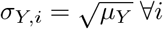, then substituting into (2) we obtain

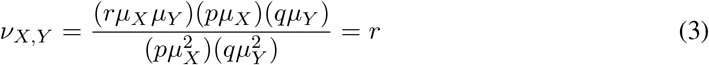

Therefore, the *ν*_*X,Y*_ is precisely the number of matched singular vectors, as expected.

A simple, continuous relaxation of this model keeps **X** the same but modifies **Y** such that 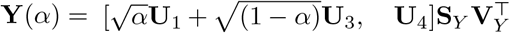, where again all **U**_i_ are orthonormal and **U**_*i*_ ⊥ **U**_*j*_ for *i* ≠ *j*, and *α* ∈ [0, 1]. From this model we obtain (see SI for detailed derivation)

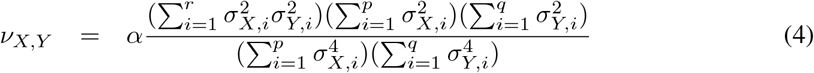

which is simply (2) multiplied by *α*. Therefore, in the case where the dimensionalities of **X** and **Y** are maximized then we have, *ν*(**X, Y**(*α*)) = *αr*. This makes clear that the ENSD reports the expected degree to which the matched singular vectors represent the basis for each dataset. Notably, *γ*_*X*_ and *γ*_*Y*_ are constant with respect to *α*, so *τ*_*XY*_ ∝ *αr* as well and achieves its upper bound 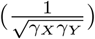 when *α* = 1.

### The ENSD captures directions of shared variability

So far we have considered the case of flat eigenspectra, meaning that the variance explained in each direction defined by the set of eigenvectors was equal. Therefore, the variance of **X** and **Y** would either completely overlap or be contained in the nullspace of the other matrix, and thus not contribute to the ENSD. In general, however, the variance of the data defined by a matrix will not be evenly distributed among the principal axes. That is, for an *n* × *p* matrix **X**, we will tend to have *γ*_*X*_ somewhat less than *p*. Notably, if there is any decay in either of the spectra of **X** or **Y**, then for the example in (2) we should observe *ν*_*X,Y*_ < *r*, reflecting the fact that *ν*_*X,Y*_ is the *effective* number of shared dimensions and not the *strict* number of shared dimensions. By “effective” we mean that not all aligned dimensions are utilized equally and that contributions by relatively low-variance modes may be neglected in the reckoning of shared dimensionality.

We can see this by inspecting equation (1). Consider two *n* × *p* matrices, **X** = **US**_*X*_ **V** ^⊤^ and **Y** = **US**_*Y*_ **V** ^⊤^, which are identical except for the concentration of their spectra. These are defined by the normalized singular values 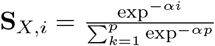 and 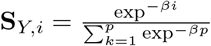. The decay parameters {*α, β*} are therefore measures of the concentration of the spectra. Suppose we fix *α* > 0 to some finite value, and vary *β*. As *β* increases, the spectrum of **Y** decays more quickly and *γ*_*Y*_ decreases (Fig.2A). Importantly, this decrease is not linear - the derivative of *γ*_*Y*_ with respect to *β* is the sum of exponentials with different decay rates (see SI). This results in an initial rapid decrease in *γ*_*Y*_ towards a lower limit of 1. With respect to *τ*_*X,Y*_, we find that the similarity of the two matrices increases with *β* (Fig.2B). As *β* increases, the overlap between basis vectors corresponding to eigenvalues of larger magnitude increases. The maximum value of *τ* is achieved when *β* and the eigenspectra of *Y* is concentrated on the first eigenvalue. In this case, 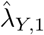 is 1 and all other 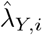 are zeros. The matrix **Λ** is thus all zeros except for the first entry, which is equal to 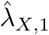. Since **W** is the identity,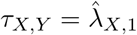.

**Figure 2:**
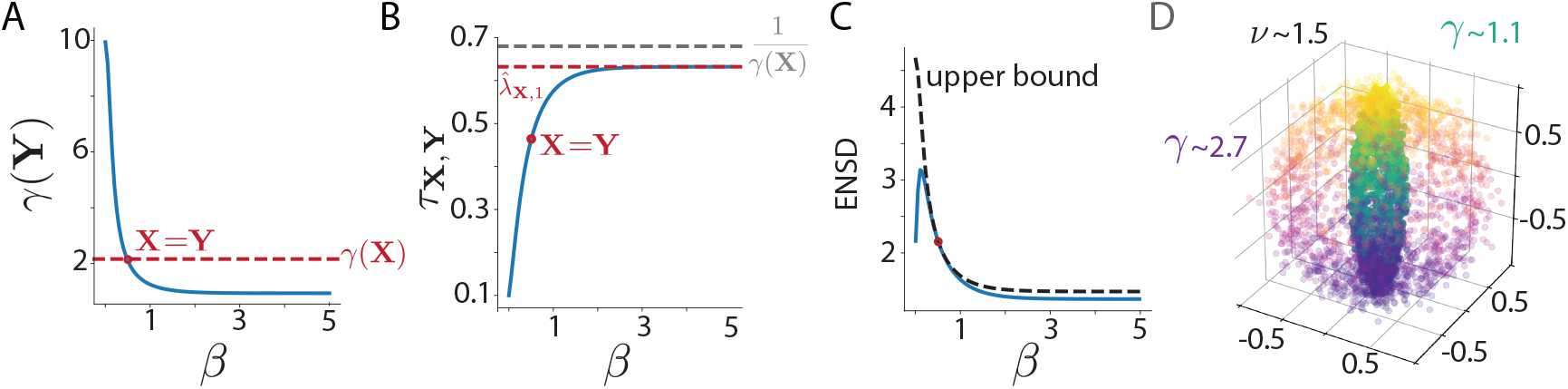
Behavior as a function of the spectrum decay rate *β* of **A**. the participation ratio **B**. similarity measure *τ* and **C**. ENSD. **D**. Toy data showing amplification of shared dimension through *ν*.

In this limit, the shared dimensionality is contained entirely in the first principle component of **Y**, and *ν*_*X,Y*_ → 1. For lower values of *β*, the ENSD peaks and then decreases (Fig.2C). Only when **X** = **Y** does *ν*_*X,Y*_ achieve its upper bound, although interestingly this is not the maximal value *ν*_*X,Y*_ achieves. Indeed, and perhaps counter-intuitively, *ν*_*X,Y*_ can be larger than the smaller of the two participation ratios.

To understand why this happens, consider *τ*_*X,Y*_ again. The outer product of the two eigenspectra, **Λ**, ends up being larger than the smaller of the eigenspectra, thus contributing dimensions of variability to *τ*_*X,Y*_ that are not captured by the *γ*_*X*_. Put more intuitively, the variability in the latter dimensions of **Y** effectively amplifies the variability in the directions of **X** that make a negligible contribution to *γ*_*X*_.

For a more concrete example of this mode amplification phenomenon, consider data lying on the surface of a sphere generated by uniformly sampling spherical coordinates *ϕ* and ψ (Fig.2D). The PR of a representative sample of this data is ∼ 2.7. If we scale the data to shrink within the x-y plane, the PR decreases to a value closer to ∼ 1.1, as the variability is mostly distributed along the z axis. However, if we calculate the ENSD between the sphere and the corresponding ellipsoid, we see that the shared dimensionality is larger, at ∼ 1.5. Thus the ENSD more accurately captures the true shared structure in the two datasets along the shared *x* and *y* axes via the amplification described above.

## 4 Comparison to existing methods

### Analysis of dimensionality estimation

In this section we demonstrate how the properties of ENSD for dimensionality estimation relate to those of two alternative methods, based on CCA [17] and reduced rank regression (RRR) [18]. For estimating shared dimensionality using CCA, we estimate dimensionality by taking the number of significant (*p*<.05) cannonical correlations. While we are aware that many variants of CCA exist [41], we focus on the classical CCA of Hotelling [17]. For dimensionality estimation using RRR we used the cross validation procedures outlined in [33]. We generated data using the probabilistic CCA model [3, 5] with two data sources each with observation dimension 30, 5 shared latent dimensions and private latent dimensions of 10 and 15 (details in the SI). In addition to noise, we randomly sparsified the observations at two levels.

Two observations are apparent from our analysis (Fig.3A). First, the ENSD is the most robust of the three methods to variations in both sample size and observation sparsity. Second, the ENSD is systematically biased downwards from the number of shared dimensions used to generate the data. This bias is consistent with the interpretation of ENSD as the *effective* number of shared dimensions and not the *strict* number of shared dimensions. As we discussed in Section 2, any degree to which there is unequal variance of shared components would contribute to a deviation in the estimated number of dimensions. We therefore expect ENSD to routinely report shared dimesionsionality that is somewhat smaller than the strict number of shared dimensions.

**Figure 3:**
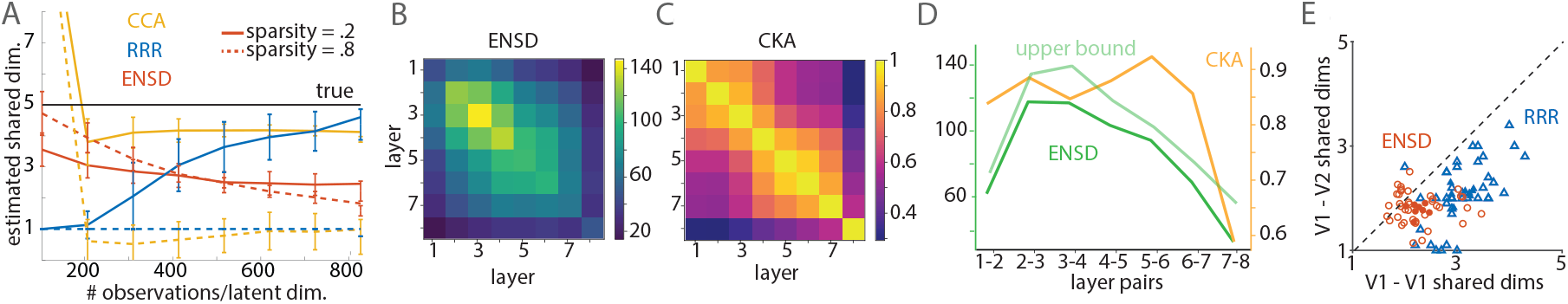
**A**. Comparison of dimensionality estimation accuracy for 3 methods in simulated experiment, averaged over 50 experiments at different sample sizes and two levels of sparsity. **B**. Across layer ENSD and **C**. CKA averaged over 6 networks. **D**. Average ENSD, upper bound and CKA across layers (off-diagonals of B and C). **E**. Comparison of shared dimensionality estimates between visual cortical areas using RRR [33] and our method. Each open marker represents a stimulus orientation for a given session. Closed markers are averages over sessions.

### Application of ENSD to artificial neural networks

Inspired by a recent study [20] which explored representational similarity within and between layers of convolutional neural networks (CNNs) using CKA, we applied the ENSD to measure the shared dimensionality between layers of a CNN. We trained a *T*in*yTen* CNN [35] to classify the CIFAR-10 dataset and computed both the CKA and ENSD between all pairs of layers, repeating this across six random initializations of the same network. The average ENSDs and CKAs across initializations are shown in (Fig.3B,C). This analysis reveals that the dimensionality of representations as measured by the PR (main diagonal, fig.3B) first increases and then decreases with layer depth. The ENSD between subsequent layers (eg. along the 1st diagonal) follows this same pattern of expansion and contraction (Fig.3D). This kind of initial expanding of dimensionality may be helpful in expanding representations for ease of learning classification boundaries.

In contrast, the average between-layer CKA remains relatively constant across layers (Fig.3C). We note that the CKA can be interpreted as the *fraction* of the maximum possible ENSD that the measured system uses (see section 2). That the ENSD includes dimensionality information therefore provides a more detailed assessment of the relationships between representations within a network.

## 5 Application to neural datasets

### Re-analysis of visual electrophysiology data

In a recent study, Semedo et al. [33] examined correlated neural activity between primary and secondary visual cortices (i.e. V1, V2). Their analysis demonstrated that correlations in trial-by-trial fluctuations were detectable both within and between regions and that these correlations were indicative of a shared (“communication”) subspace that displayed dimensionality similar to the dimensionality of the stimulus used in the task. The authors used RRR for determining the dimensionality of this subspace. We propose that our measure can be equivalently used to conduct this analysis without model fitting.

We repeated their analysis and applied the ENSD to their data (Fig.3E). We obtained similar dimensionality for both target V1 and V2 populations from a size-matched source V1 population, although our results are somewhat downward-biased, as expected (see Fig.3A). Because we were able to conduct our analysis without model fitting, computation time was an order of magnitude smaller using ENSD as compared to RRR (see SI), opening the door to analysis of much larger-scale datasets in the future.

### Analysis of olfactory connectivity data

The relationship between structure and function in the olfactory system of the fruit fly, *Drosophila Melanogaster* has been studied in increasing depth and detail over recent decades. Significant progress has been made in revealing the computational logic of the fly*’*s olfactory circuits as a result of this focus. Here, we demonstrate that an ENSD based analysis reconfirms a number of important findings across data modalities and experimental paradigms, and affords novel observations, which open up avenues for further research.

In the fly, chemical signals in the environment (odorants) evoke activity in 51 genetically defined channels (glomeruli), with different odorants activating different subsets of these channels. While these channels vary in the breadth of their tuning to odorant features [16], a primary odorant-type tuning has been assigned for each channel. For example, a channel may be classified as primarily sensitive to pheremones or food sources e.g. decayed fruit (Fig.4F) [7, 39]. Channel activations are relayed via projection neurons (PNs) in the antennal lobe (AL) to two downstream areas (Fig.4A): i) the mushroom body (MB), an area that has been shown to mediate adaptive olfactory learning [2] and ii) the lateral horn (LH), a relatively poorly characterized area that is thought to mediate innate olfactory behaviour [9]. Over the past decade, these circuits have been the focus of intense research attempting to understand how information is represented across these channels [4, 6, 39, 31, 7, 22], both in an attempt to understand which odorant features are relevant to olfactory processing as well as, more generally, how biological systems represent complex, high-dimensional environments without any clear statistical structure.

**Figure 4:**
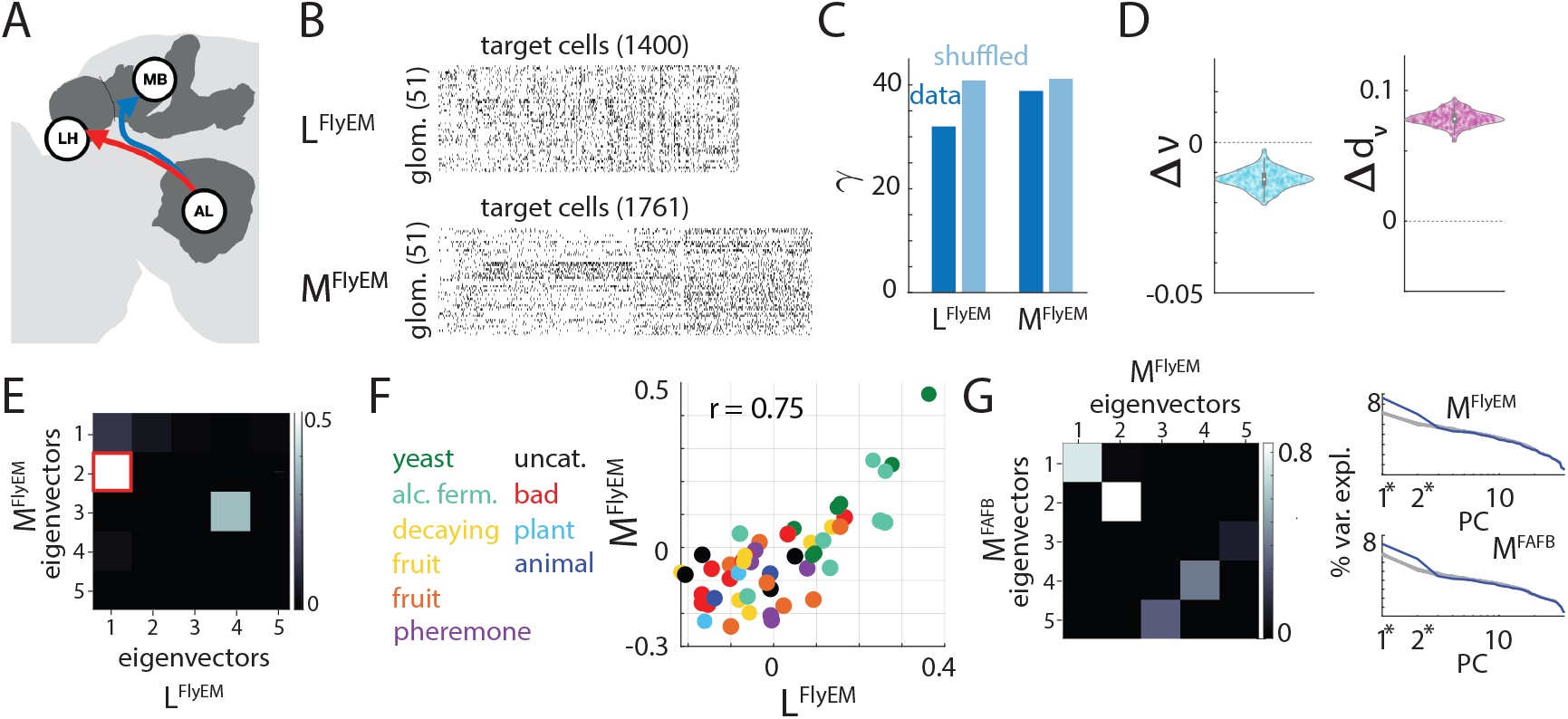
**A**. Schematic of the fly olfactory system showing parallel pathways in the lateral horn (LH) and mushroom body (MB) which both recieve input from the antennal lobe (AL). **B**. Binarized connectivity matrix between the 51 types of glomeruli in the AL and the principal neurons in the LH or MB. **C**. PRs of the connectivity matrices for the LH (left) and MB (right) versus a shuffled null models. **D**. Effect of shuffling on the ENSD (left) and the distance (right) between the LH and MB. **E. W** matrix of eigenvector overlaps between the LH and MB. Eigenvector overlaps corresponding to the top 5 eigenvalues shown, all statistically insignificant overlaps (*p* > 0.05), are set to zero. **F**. Loadings of the different glomeruli onto the eigenvectors found in E. (right), color-coded by their preferred odorant type (left). **G. W** matrix showing eigenvector overlaps between the two MB datasets (left) and their corresponding eigenspectra (right) compared to null models (grey line).

To probe the logic of governing inputs to these downsteam areas, we turn to two recently released connectivity datasets: the Hemibrain connectome, which contains a detailed description of inputs to LH and MB from a single fly (matrices **L**^*FlyEM*^ and **M**^*FlyEM*^ respectively, Fig.4B), and the FAFB dataset, which describes input to a smaller collection of MB principal cells from a second individual (**M**^*FAFB*^) (for data pre-processing see SI) [30, 39]. For each of the three empirical binary connectivity matrices, we define a corresponding ensemble of randomized matrices, 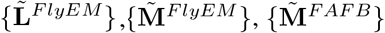. Our randomizing procedure removes correlations between channels but preserves the marginal statistics of the data matrices [6] (see SI). These three datasets allow us to compare shared structure within and across individuals.

A key question regarding these circuits is the extent to which the feedforward connectivity contains interpretable structure. This can be achieved by computing the participation ratio of these data and comparing these values to the corresponding null models [6]; this analysis reveals that i) all three connectivity matrices contain a degree of nonrandom organization and ii) **L**^*FlyEM*^ is significantly more structured than either **M**^*FlyEM*^ or **M**^*FAFB*^ [6, 39] (Fig.4C, the difference in d_*ν*_ and *ν* between data and shuffled connectivity matrices is significantly larger, analysis of **M**^*FAFB*^ in the SI). This latter point is also consistent with the behavior hypothesized to be mediated by these areas: an associative learning center should be higher dimensional in order to increase memory capacity [22], whereas mediating innate behavior likely requires more structure and should therefore be lower dimensional.

To determine if the structure identified in the two pathways is related, we can use the ENSD and its derivatives to explore the shared input subspace. Comparing the shared subspace to that of shuffled data will allow us to determine whether any observed structure is significant or simply the expected result of the marginal statistics of the data. To this end, we compute *ν*(**L, M**) and d_*ν*_ (**L, M**) for the *FlyEM* dataset and compare these statistics to the statistics obtained when the MB dataset is shuffled. These analyses reveal that, compared to what would be expected for fully random MB connectivity, the LH shares a representational subspace of higher dimensionality with the MB, corresponding to more similar input connectivity patterns (Fig.4D). This suggests a novel result: that inputs to the MB and LH are partially aligned.

To probe this alignment further, we can look at overlaps between pairs of high variance eigenvectors via the **W** matrix. Our analysis reveals a single statistically significant and strongly overlapping dimension, the first principal component of **M**^*FlyEM*^ and the second PC of **L**^*FlyEM*^ (Fig.4E). The strongly aligned subspace is therefore 1-dimensional. We can interpret this shared dimension by comparing eigenvector loadings, which reveals an organization dominated by a set of channels tuned to food, specifically yeast and fermented fruit related odorants (Fig.4F). This is consistent with findings from [7], however our analysis also demonstrates this food dominated shared subspace is statistically significant i.e. not simply a product of the marginal statistics of the data, and that it is consistent across individuals (see SI). Each of these results is consistent within (*FlyEM* dataset) and across individuals (*FlyEM*-*FAFB*, see SI). The food-tuned shared subspace between LH and MB inputs revealed by our ENSD based analysis may play an important role in the circuitry that permits olfactory learning in the MB pathway to alter innate food related behaviour (LH pathway), as recently observed in [12, 21] - this could be explored in further computational studies. Overall these analyses reveal that part of the structure observed in inputs to the LH and MB is shared between the two areas and that this shared subspace relates to an ethologically relevant odor scene.

Across-animal analysis confirms that the first PCs of the two MB datasets are aligned and food tuned [39]. Examination of the **W** matrix indicates that the second PC is also aligned (Fig.4G, left). As PCA showed that only the first two dimensions of these matrices differ from the null model (Fig.4G, right), we conclude that all of the structure in the MB inputs is shared across the two individuals. This supports the hypothesis that the small structured component in MB inputs is genetically determined rather than learned.

### Further analyses in the SI

To complement this analysis, in the SI we use the ENSD and its derivatives to examine the shared variability between datasets of two different modalities: odorant evoked neural activity in PNs and the corresponding PN connectivity patterns. We find that inputs to the LH, but not the MB, are significantly tuned to the statistics of channel activation. In the SI also we show that the above results are robust across choice of synaptic threshold for the FlyEM dataset and that marginal statistics of the connectivity data are not sufficient to account for the consistent structure observed.

## 6 Discussion

Here we have presented the ENSD, a generalization of the participation ratio to measure the dimensionality of shared subspaces in paired uni or multimodal datasets. We have shown that the ENSD has a number of advantages compared to existing, model-based methods for estimating shared variability: it is robust to sample size and observation sparsity, requires minimal computation time (just 5 matrix multiplications), and its component parts are easily interpretable. The ENSD also reduces to the CKA score through a scaling factor and offers an interpretation of this score as a fraction of the possible shared dimensionality that a system is using. Because of the relationship to CKA, we expect that critiques of that measure likely also apply to the ENSD, including a lack of sensitivity to low variance dimensions [10, 11, 26].

Another important aspect to consider is the robustness of our shared dimensionality estimator to noise. A previous analysis of linear dimensionality estimation revealed that the PR tends to overestimate the embedding dimensionality when significant noise is added to synthetic data and when the underlying data manifold is nonlinear [1]. However, amongst common shared dimensionality estimators, the ENSD changes the least in response to variations in sample size as well as in response to changes in random sparsening rate (Fig3.A). Both of these features are indicative of a relative robustness to various sources of noise. In contrast, CCA appears to be particularly sensitive to both small sample sizes and high sparsity rate, and RRR seems similarly sensitive to sparsity. These results indicate that RRR and CCA would both be poor candidates for analyzing the highly sparse connectivity data presented in Section 5. Moreover, CCA and RRR both require multiple hypothesis testing to estimate shared dimensionality, a requirement that is troublesome for exploratory analyses.

As demonstration of the utility of the ENSD, we applied it to two neural data sets of different modalities and data types (dense vs sparse). We were able to recover existing results from electrophysiology experiments that probed communication subspaces between V1 and V2 [33], and from analyses of olfactory connectivity that showed a subtle food-related structural component in the inputs to MB [6, 39]. Probing the ENSD of the fly connectivity matrices strategically also revealed consistent and interpretable shared structure between inputs to LH and MB pathways of the fly olfactory system within and across individuals, refining the findings in [7] as well as providing new insights into the potential genetic origin of this shared structure. That we were able to recover all of this from application of a single measure to connectomic data serves to showcase the power and interpretability of the ENSD. The same approach can easily be applied to connectomic data to compare structure across other individuals when further data is released [14] or combined with electrophysiology data (see SI for early experiments in multimodal analysis).

The ENSD offers a simple and interpretable method for analysing shared variance between multiple paired high-dimensional data sets. We have demonstrated use cases in neuroscience and machine learning contexts, but this technique can equally be applied in other fields, from physics to sociology, to ask related questions about shared variance contained in data. Our future directions include exploring how the ENSD can be applied to time-lagged neural responses to characterize e.g. representational drift [32], and what the ENSD can tell us about representations across neural network architectures. Additionally, recent work has incorporated scale-dependence in the PR, which can reveal differences in local and global structure that are be obscured by other dimensionality measures [27], with application to spatiotemporal data. The ENSD can be easily extended and applied in a similar manner. We are also interested in extending the measure to non-linear dimensionality metrics via kernelization of the ENSD.

## Supplementary Information: Effective number of shared dimensions between paired datasets

### 1 Properties of *ν*_*X,Y*_

For the following examples, consider the *n* × *p* matrix **X**, and the *n* × *q* matrix **Y**, where *n* > *p, q*.

#### 1.1 Equality with *γ*_*X*_

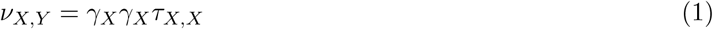

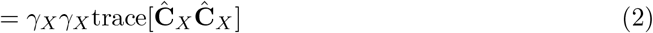

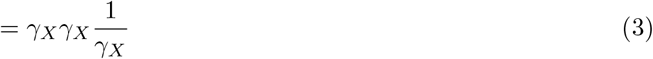

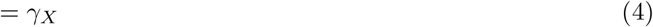

##### 1.2 Upper bound

Just as *γ*_*X*_ is upper bounded by *p*, the ENSD also admits an upper bound. To see this, first recall that the Cauchy-Schwartz inequality gives

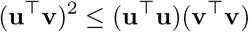

for any vectors **u** and **v**. Noting that **u** and **v** in the above expression may be the vectorizations of matrices **U** and **V** we may equivalently write

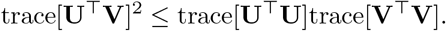

Substituting **U** = **Ĉ** _*X*_ and **V** = **Ĉ** _*Y*_ gives

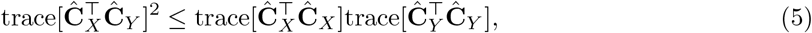

and using the notation presented in Section 2 of the main paper we may write inequality (5) as

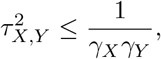

and therefore

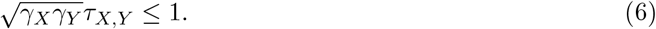

Finally, multiplying both sides of inequality (6) by 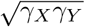 gives the upper bound for *ν*_*X,Y*_,

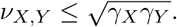

and additionally, it is clear that *τ*_*X,Y*_ is upper bounded by the inverse of the geometric mean of the participation ratios, 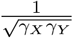.

##### 1.3 Decomposition in terms of eigenvalues and eigenvectors

The ENSD may be rewritten in terms of the eigenvalues and eigenvectors of it*’*s constituent matrices, via the scaled diagonalized covariance matrix, trace[**Ĉ** _*X*_] = **U**_*X*_L_*X*_ **U**_*X*_, where L_*X*_ is a matrix with eigenvalues 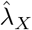 on the diagonal and **U** are the principal axes of the data. We first rewrite the trace term,

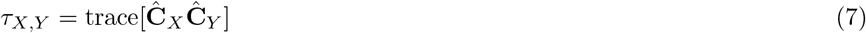

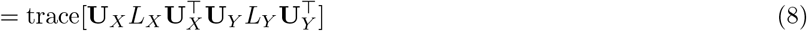

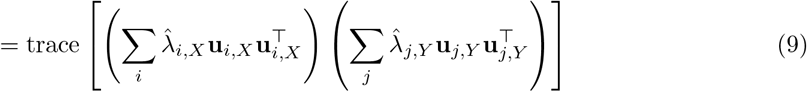

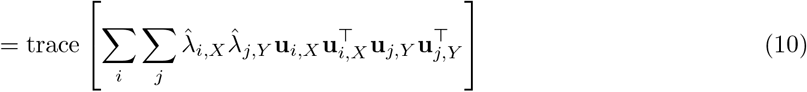

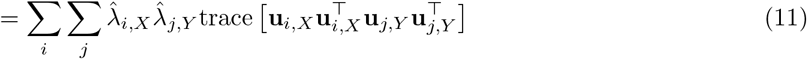

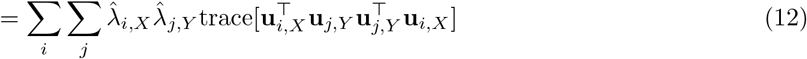

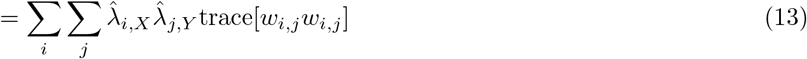

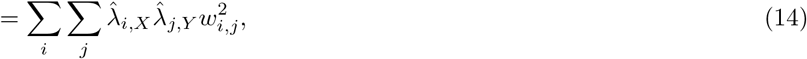

where 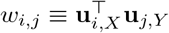. Equation (14) explicitly shows how *ν*_*X,Y*_ is a function of both the eigenvalues of **Ĉ** _*X*_ and **Ĉ** _Y_ and also the inner products of their constituent eigenvectors. If we rewrite the participation ratio in the same way, we find that w_*i,j*_ =1 ∀*i, j*, so

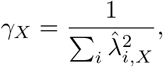

allowing us to rewrite the ENSD as

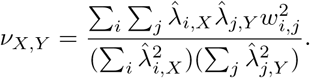

Since every term in (14) is indexed by *i* and *j* we can rewrite each in terms of the *i, j*^*th*^ entries of a matrix. Let 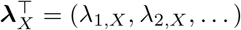, and

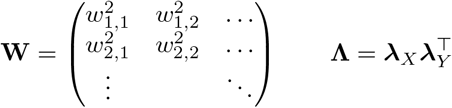

then

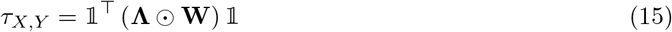

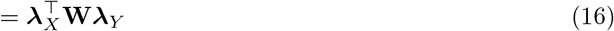

and

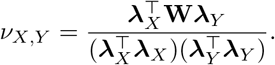

This way of writing out *τ*_*X,Y*_ allows for an intuitive interpretation of this term: it captures the overlap between two subspaces with **W**, which is then scaled by the respective eigenspectra, **Λ**. This similarity measure is then weighted by the individual dimensionalities of the two subspaces in the equation for *ν*_*X,Y*_ to arrive at a final numerical estimate for the dimensionality of the shared subspace.

The corresponding distance metric, according to [8], may also be expressed in this way, as

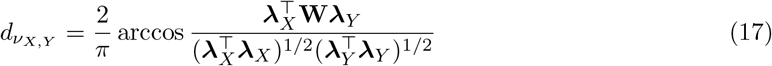

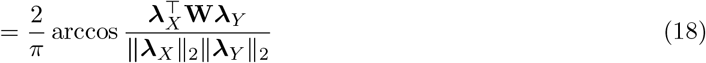

### 2 Probing the ENSD using synthetic data

#### 2.1 Partially shared subspaces

For the simplest toy example we construct a scenario where we define the number of shared dimensions between **X** and **Y** to be an integer *r*. Suppose the singular value decompositions of these matrices are given by the *n* × *p* matrix 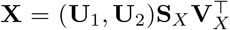, where **S**_*X*_ = diag(*σ*_*X*,1_,. .., *σ*_*X,p*_) and the *n* × *q* matrix 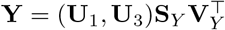, where **S**_Y_ = diag(*σ*_*Y*,1_,. .., *σ*_*Y,q*_), where the key feature we impose is that **U**_i_ ⊥ **U**_j_ for *i* ≠ *j*. The shared subspace is defined by the *n* × *r* matrix **U**_1_, while **U**_2_ is *n* × (*p* − *r*) and **U**_3_ is *n* × (*q* −*r*).

The PR for **X** can be written as

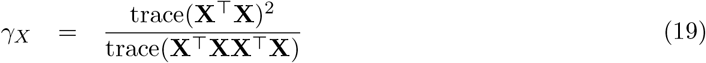

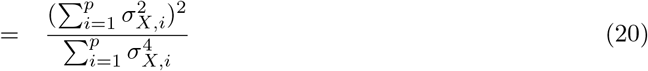

and *ν*_*X,Y*_ can be written

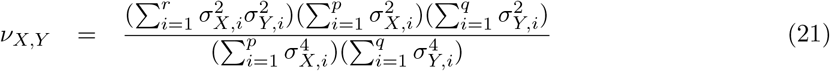

or, as described in the main text, if the dimensionality of both matrices are maximized (corresponding to flat eigenspectra),*ν*_*X,Y*_ reduces to exactly the number of shared eigenvectors.

#### 2.2 Continuous partially shared subspaces

An illustrative extension of this simple example is a continuous relaxation of this model with the modification that 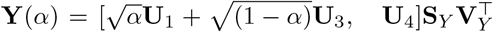, where again all **U**_*i*_ are orthonormal and **U**_*i*_ ⊥ **U**_*j*_ for *i* ≠ *j*, and *α*∈ [0, 1].

We then have

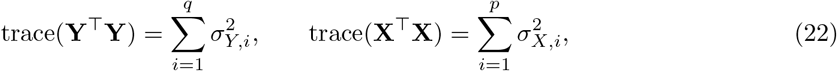

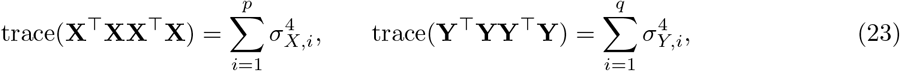

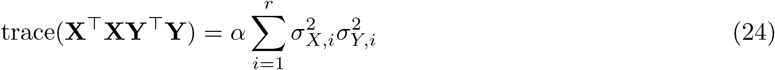

Therefore,

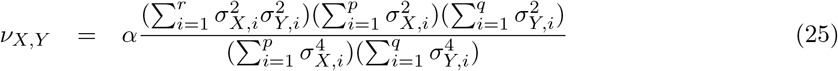

We can see that (25) is simply (21) multiplied by *α*. Therefore, in the case where the dimensionalities of **X** and **Y** are maximized (i.e. 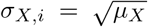 and 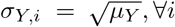) then we have, as expected, *ν*(**X, Y**(*α*)) = *αr*. Additionally, since *γ*_*X*_ and *γ*_*Y*_ are constant, *τ*_*XY*_ ∝ *αr* as well and achieves the upper bound 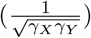 when *α* = 1.

**Supplementary Figure 1:**
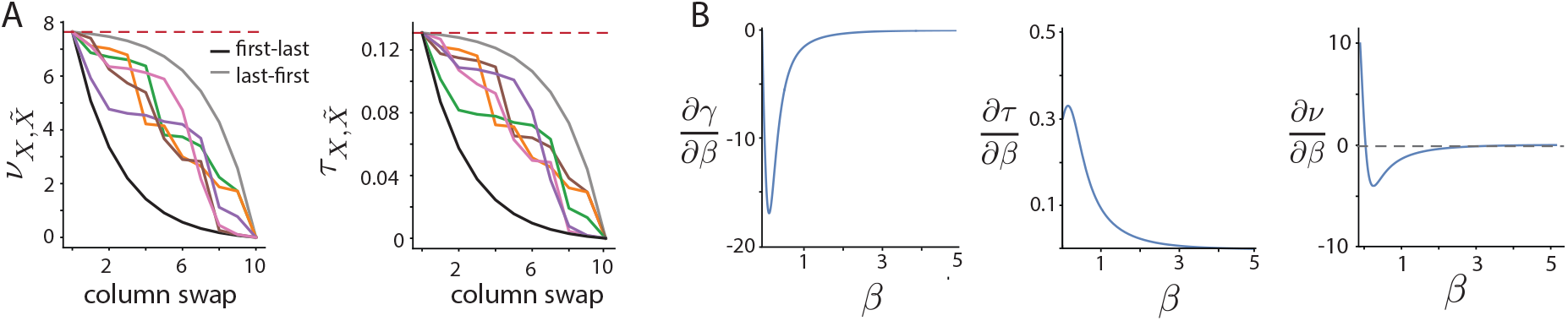
**A**. Plot of 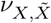 and 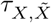 when swapping out *n* columns of U to generate 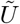 in a particular order – the column corresponding to the first eigenvalue to last (black line), last to first (grey line) or in random order (colored lines). **B**. Plots of the derivatives of the PR (*γ*, left), the similarity term (*τ*, center) and the ENSD (*ν*, right) as a function of the eigenvalue decay of matrix **Y**.

#### 2.3 Dimensions explaining more variability contribute more to *ν*

Consider an *n* × *p* matrix 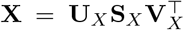, where **S**_*X*_ = diag(*σ*_1_,. .., *σ*_*p*_) and *σ*_i_ > *σ*_i+1_, and a second basis 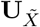, where again all columns of **U**_i_ are orthonormal and **U**_i_ ⊥ **U**_*j*_ for *i* ≠ *j*. We now generate a second matrix, 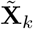, which is identical to **X** *except* that the kth column of **U**_*X*_ is replaced with the kth column of 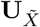. Since all columns of **U**_*X*_ and 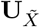 are mutually orthogonal, 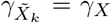 irrespective of which column of **U**_*X*_ is swapped. The only term in *ν*_*X,X*_ that is affected is the similarity term, 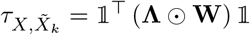. Since the matrices are identical except for their eigenvectors, swapping columns of **U**_*X*_ only affects the matrix of eigenvector overlaps, **W**.

For *i, j* ≠ k, W_*i,j*_ = 0, as the associated eigenvectors are orthogonal, and for all *i* = *j* ≠ k, W_*i,j*_ = 1, as 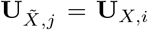. When *i, j* = k, now W_*k,k*_ = 0, whereas before the swap it was equal to 1. Since the kth term in the sum (14) is now zero, 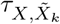 decreases by an amount, 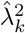. The eigenvectors are strictly decreasing, so swapping out the kth eigenvector always results in a larger change to 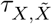, and therefore 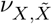, than swapping out the (k + 1)th vector.

An example of this is shown in figure 1A, where we plot the ENSD and 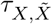 as a function of the number of columns swapped in 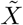 for different swapping orders. In this case, both **X** and 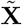 are of dimensionality ∼ 7.5. In the case where we successively swap the columns in **U**_*X*_ for columns in 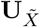 from largest to smallest eigenvalues (black line), 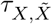 and 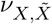 decrease more quickly than if the columns were swapped from smallest to largest eigenvalue (grey line). Random column swaps (colored lines) all produce 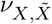 and 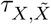 within the envelope defined by these two modifications.

#### 2.4 The ENSD captures amplification of shared variability

Finally, in section 3 of the main text we consider the case where the eigenspectra of the two matrices have different decay rates which produces some interesting phenomena. We consider two *n* × *p* matrices, **X** = **US**_*X*_ **V**^⊤^ and **Y** = **US**_*Y*_ **V**^⊤^, so **X** and **Y** are identical except for their rates of eigenvalue decay. The decrease in the PR is not linear, as shown by the plot of its derivative with respect to *β* (Fig.1B, left). Likewise, the derivative of *τ*_*X,Y*_ is strictly positive but decreasing over time as *τ*_*X,Y*_ approaches the upper bound (Fig.1B, center). In this limit, the shared dimensionality is contained entirely in the first dimension of **X** and **Y**, and *ν*_*X,Y*_ → 1. The ENSD, however, shows some interesting behavior, first increasing then decreasing (see Fig.2C in the main text). We might expect that the ENSD would strictly decrease as the matrices become more different. However, its derivative goes from positive to negative in the brief period that 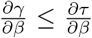 (Fig.1B, right), which causes the shared dimensionality to be greater than the participation ratio. After the inflection point, 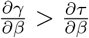 the shared dimensionality is again below the participation ratio, and the decrease in ENSD is mainly due to the decrease in participation ratio of **Y**.

## 3 Comparison of shared dimensionality estimation techniques

We sought to evaluate the ENSD compared to alternative measures of shared dimensionality using synthetic data where ground truth is known. To our knowledge, only two alternatives exist: model-based estimation of shared dimensionality using reduced rank regression (RRR) and statistically-based estimation using canonical correlation analysis (CCA). For the RRR method we followed the procedure outlined in [7]. Briefly, the RRR model is used to predict variability in one dataset from the variability of the other. Models are constructed with dimensionalities varying from 1 to k and each assessed via cross-validation. The model dimensionality with the best predictive CV score is selected as the shared dimensionality. For the CCA method, we conducted CCA using canoncorr() function in Matlab (version 9.13.0. Natick, Massachusetts: The MathWorks Inc., 2022.). Each canonical correlation with a p-value of <0.05 (corrected for false discovery rate) was considered significant. The significant canonical correlations were then summed to give the estimate of dimensionality.

**Supplementary Figure 2:**
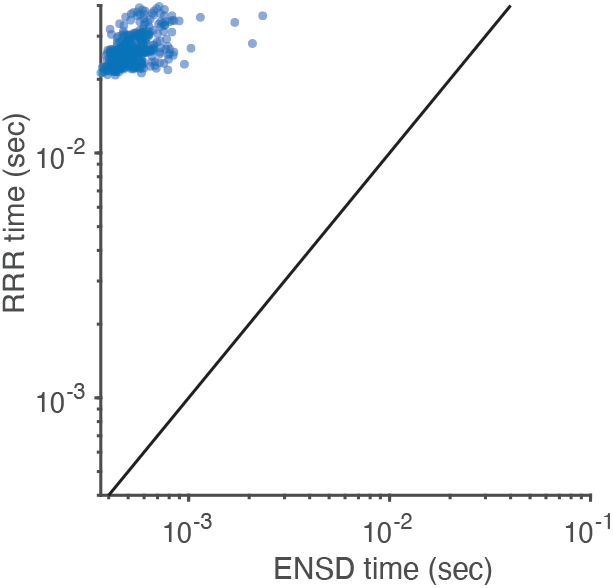
Wallclock time of electrophysiology data analysis using ENSD and RRR. Each marker represents a single run of the analysis.

Synthetic data was generated using the probabilistic CCA (pCCA) model [1, 3]. Briefly, the pCCA model is a structured factor analytic model where in observations of vector measurements for two data sources (**y**_A_, **y**_B_) are determined by P latent variables whereby

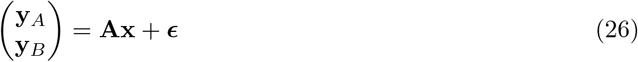

where ϵ **∼** 𝒩 (0, **D**), with **D** diagonal, and **x ∼** 𝒩 (0, 𝕀_*P*_). The shared subspace is structured through the partitioning of **x** such that 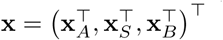 where **x**_*A*_ is latent variability associated exclusively with measurements **y**_*A*_, **x**_*B*_ is latent variability associated exclusively with measurements **y**_*B*_, and **x**_*S*_ is latent variability associated with both measurements **y**_*A*_ and **y**_*B*_. The latent dimensionality *P* is given by the sum of the dimensionality of all components; *P* = *P*_*A*_ + *P*_*S*_ + *P*_*B*_. The mixing matrix is structured to maintain the partitioning of variability via the following block structure

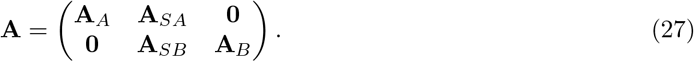

In addition to the noise variable **ϵ**, we tested the robustness of these methods by randomly setting each observation to 0 with 2 different probabilities (0.2 and 0.8). We simulated data using this model with 31 observations of each **y**_*A*_ and **y**_*B*_, *P*_*A*_ = 10, *P*_*B*_ = 15, *P*_*S*_ = 5. Sample sizes ranged from 50 to 500 and each sample size was repeated across 50 different experiments. Nonzero elements of **A** were drawn iid from a standard normal distribution for each experiment. Means and standard deviations reported in Figure 3 are taken over all experiments.

## 4 Re-analysis of visual cortex electrophysiology data

For the electrophysiology data analysis in Section 5, we conducted estimation of dimensionliaty via RRR according to [7] using code retrieved from the authors*’* github repository https://github.com/joao-semedo/communication-subspace. The data was retrieved from a publicly available CRCNS repository [10]. Data collection was described in detail in [9].

Briefly, an anesthetized macaque was presented with a visual stimuli consisting of a series of oriented gratings with one of 8 different orientations. Extra-cellular electrophysiological recordings were conducted simultaneously in V1 (97 single units via Utah array) and V2 (31 single units via tetrodes) during stimulus presentation. Each stimulus orientation was analyzed separately. For each run of an analysis, the V1 sample sizes were matched to the V2 sample size by randomly sampling both a “target” and “source” subsample from V1. V1 dimensionality was taken to be the optimal RRR model dimensionality to predict the target V1 dataset from the source V1 dataset. Shared dimensionality was taken to be the optimal RRR model dimensionality to predict V2 neural responses using the source V1 dataset as regressors. The estimation procedure was repeated 10 times for each dataset and the results were averaged. We calculated ENSD values for each of these 10 resamplings for each stimulus orientation and results for each resampling were averaged to give 1 shared dimensionality value for each stimulus orientaion. Since the ENSD is not limited to matrices that share both dimensions, we were able to complete the analysis using the full V1 dataset. The results from this analysis reveal that size-matching may be causing an underestimation of the dimensionality of the communicaation subspace between the two areas.

Although absolute computation time for each method was fast, the wallclock computation time for ENSD was about two orders of magnitude faster than for RRR (Fig. 2), which has implications for analysis times for very large datasets.

## 5 Analysis of olfactory connectivity data

In section 5 of the main text, we explore shared structure in three olfactory datasets from *D. Melanogaster*. Each is a neural connectivity matrix obtained by electron microscopy [5, 11] describing connectivity from the fly Antennal Lobe (AL) to one of two downstream areas, the Lateral Horn (LH) and Mushroom Body (MB). Two such matrices were extracted from one individual via the Hemibrain dataset: **L**^*FlyEM*^ (dimensions 108 × 1400) and **M**^*FlyEM*^ (dimensions 108 × 1761) [6]. These data describe the number of synapses between individual AL and LH/MB cells. The final matrix was obtained from a second individual via the FAFB dataset, **M**^*FAFB*^ (dimensions 109 × 1344) and describes binary connectivity between individual AL and LH/MB celxls [11]. We contract the first dimension (uniglomerular projection neuron (uPN)), by adding together uPNs associated to the same olfactory channel (originating in the same glomerulus). This results in a consistent first dimension of size *n*_channels_ = 51 across all three datasets. A threshold is then applied to binarize the FlyEM datasets, θ ∈ [1, 3, 5, 10]. θ is set to 3 in the main text. Here, we also apply the ENSD to the synaptic count data.

As we are interested in investigating shared structure between these datasets, we compare each dataset to a null model in which correlations contained in the empirical data are removed via shuffling. Our shuffling procedure for binary connectivity matrices follows [4, 11] and results in an ensemble of null model matrices, each of which has the same the marginal statistics (row and column sums) as the empirical data. We use an ensemble of size x = 300 for the *θ* = 3 analyses and x = 100 for other *θ* values. To shuffle the synaptic data, we first apply a threshold to the data and then shuffle via the above procedure. Finally, synaptic counts are added back in randomly such that the row sums are held constant, i.e. synaptic counts from each channel/glomerulus are the same as the empirical data, but column sums are allowed to diverge. Rows in each matrix, empirical or null model, are mean centered. It is important to note that in each analysis, only one matrix is shuffled (indicated by a grey bar in the figure header). We choose to shuffle the less structured matrix of the pair, i.e. **M**^*FlyEM*^ or **M**^*FAFB*^, rather than both matrices. Shuffling the matrix with more structure, i.e. **L**^*FlyEM*^, would result in a random matrix which may be (and in our case is) more similar to the less structured matrix. This similarity is, from our perspective, artificial, as we are interested in understanding how the structured components in two datasets relate to one another, not the random components.

In Supplementary Figures 3-6, we present an extended analysis of the data from the main text. This includes pairwise analyses of the three connectivity matrices in SI-Fig.3A, with synaptic count data for the **L**^*FlyEM*^ and **M**^*FlyEM*^ replacing the binary data from the main text. In SI-Fig.3B, we show that the overlap between shared dimensions (eigenvector 1 in **L**^*FlyEM*^ and eigenvector 2 in **M**^*FlyEM*^ and **M**^*FAFB*^) is significantly increased in empirical data vs the null model. In SI-Fig.4, we confirm that the shared subspace is not an artefact of threshold choice, but a robust feature of the data by analyzing the changes in *ν* and d_*ν*_ for all synaptic and binary thresholding configurations. In In SI-Fig.5, we confirm that the shared structure (aligned 1D subspace) cannot be explained by the marginal statistics of the data alone as the loadings and proportions of synapses or connections for each information channel are relatively uncorrelated.

Because our shuffling procedure significantly affects the dimensionality of the data (shuffled matrices are higher dimensional), *ν*, which is sensitive to this dimensionality via the *γ*_*X*_*γ*_*Y*_ component, may be less indicative of shared structure than d_*ν*_. For instance in the comparison between MB inputs across individuals (final column of SI-Fig.3), *ν* increases for the null model relative to the data. In this case and in others, the change in d_*ν*_ more clearly indicates shared structure that has been disrupted by shuffling. We expect that in studies where the procedure for generating samples from the null model doesn*’*t significantly change dimensionality, *ν* would be equally useful for detecting shared structure, as well as giving an interpretable value for the size of the shared variability component.

Finally, we show the connectivity data reordered by loading on the shared dimension in SI-Fig.6. The LH consists of two major cell types: output neurons (LHONs) and local neurons (LHLNs), as well as a few minor cell types (other). By reorganizing the components of **L**^*FlyEM*^ corresponding to these cell types, we can see that both the LHON and LHLN populations contain subpopulations of cells that are strongly tuned to the food related features of the shared dimension, while the other cell types do not contain any strongly tuned neurons. We also note that, while the output neurons contain subpopulations that are both positively and negatively tuned to the shared dimension, the local neuron population only contains a subpopulation that is positively tuned to these features. This is an interesting circuit feature that we intend to study in future work. The MB contains a single primary cell type, the Kenyon Cell (KC), and is physically organized into a series of lobes. Reordering **M**^*FlyEM*^ by loading onto the shared dimension reveals that all three lobes, *α*/*β, α* ^*′*^/*β* ^*′*^ and *γ*, contain subpopulations of neurons that are both positively and negatively tuned.

Additionally, we use the ENSD to probe variability shared between a subset of connectivity data from the Hemibrain dataset (*FlyEM*) and neural activity in a subset of channels obtained from [2]. The activity data describes the responses of 37/51 channels to a set of 84 odorous stimuli, including fermentation and other food related odorants/blends. The raw data is preprocessed to a matrix of Z-scored responses **R** shown in SI-Fig.7A. We find that input connectivity to the LH is significantly tuned to features of the neural responses (including but not limited to food related features. SI-Fig.7B, first column), relative to shuffled LH connectivity, whereas the MB input (second column) is more weakly tuned to the neural activity (comparison of Δ*d*_*ν*_ plots in columns 1 and 2). We confirm that the subspace shared between LH and MB inputs is also present in the subset of connectivity data used for this analysis (last column). This difference in tuning between LH and MB inputs is consistent with the putative roles of these two neuropils in innate vs learned olfactory behaviour. This analysis demonstrates that the ENSD is a simple and useful tool for uncovering consistent relationships between datasets of different modalities.

## Code availability

Code to reproduce the main results of this work is available at https://github.com/Aoi-Lab-Projects/ENSD.

---

**Supplementary Figure 3:**
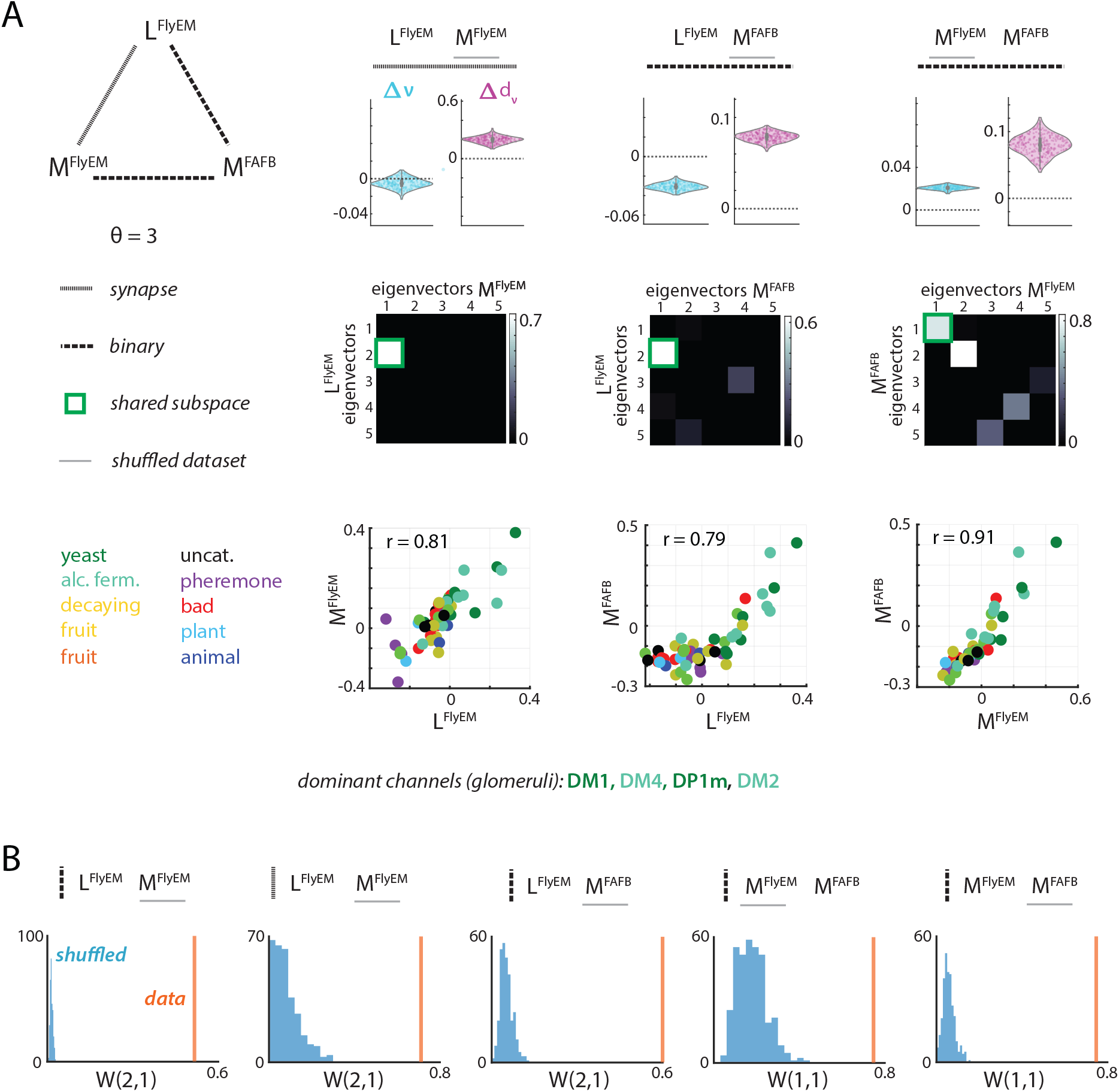
**A**. Pairwise analysis of shared structure between connectivity matrices from FlyEM and FAFB datasets. *Top row* : changes in ENSD and distance metric when one dataset is shuffled (underlined in grey). *Second row* : **W** matrices containing eigenvector overlaps corresponding to the top 5 eigenvalues shown, all statistically insignificant overlaps (*p* > 0.05), are set to zero. *Bottom row* : Comparison of loadings onto shared dimension. **B**. The shared structure identified in **W** is not preserved when one dataset is shuffled.

**Supplementary Figure 4:**
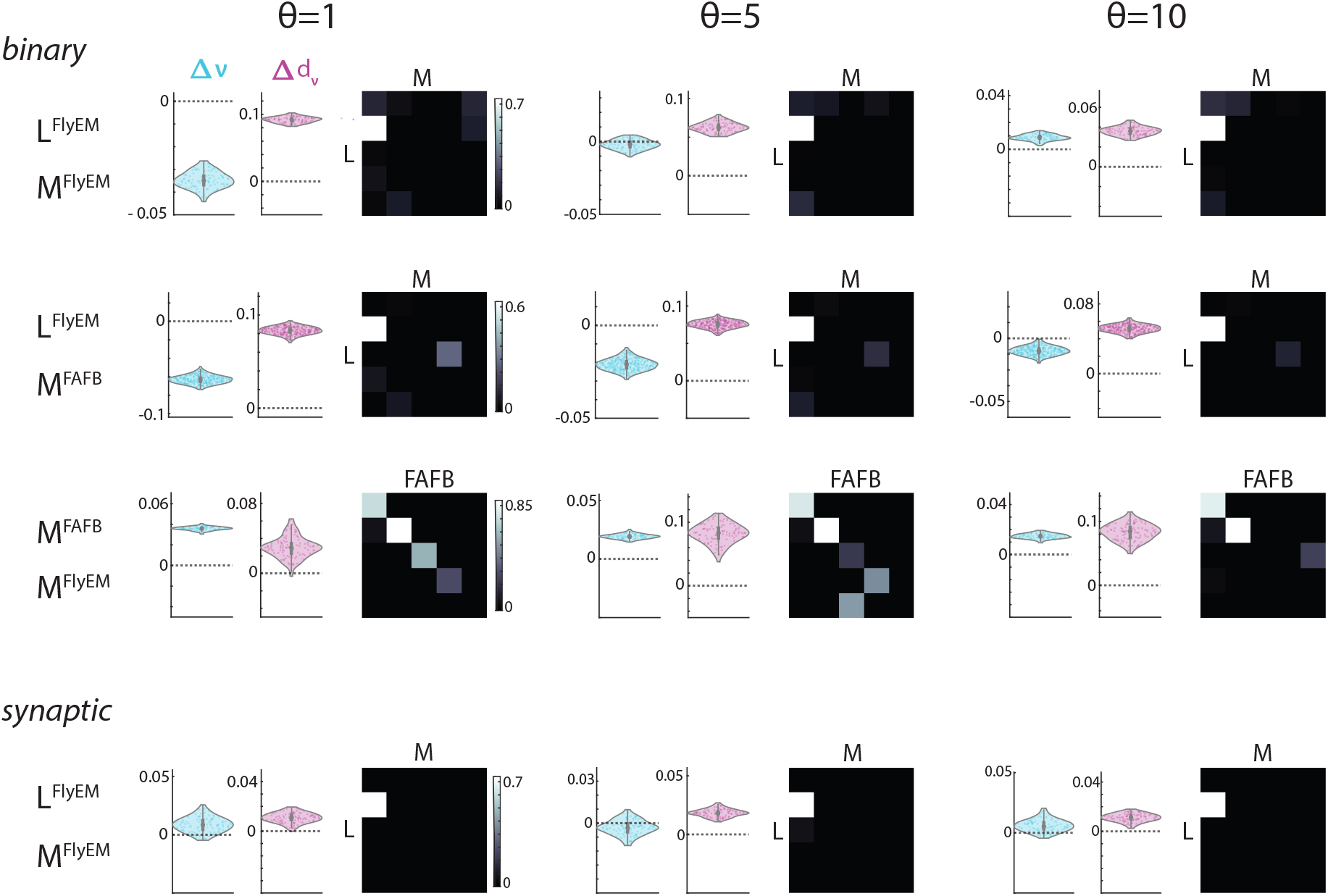
Shared subspace analysis from the main text is consistent across a range of synaptic threshold values.

**Supplementary Figure 5:**
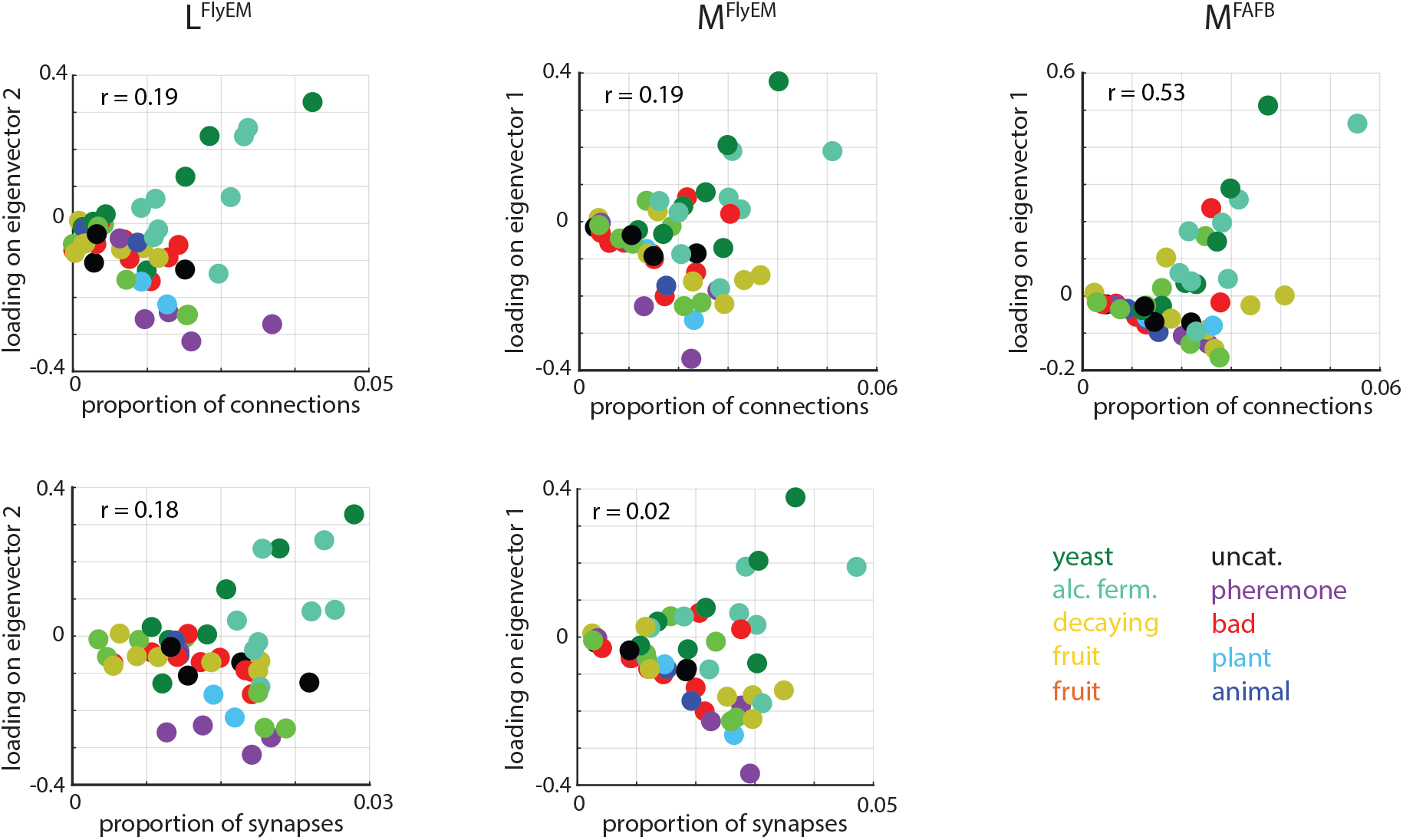
Loadings on shared dimension from each dataset are compared against proportion of connections/synapses for each olfactory channel (glomerulus).

**Supplementary Figure 6:**
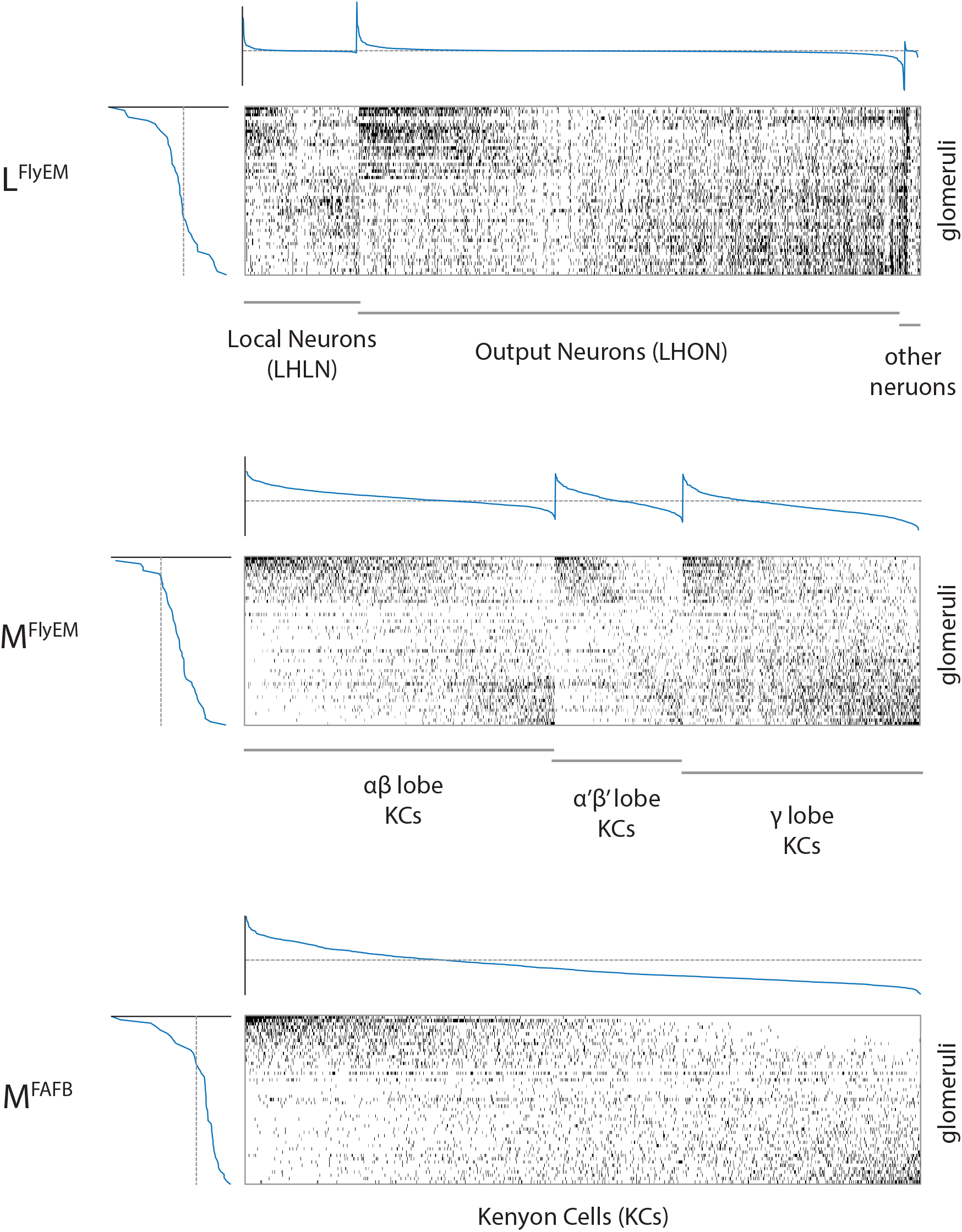
Binary connectivity matrices ordered in both dimensions by loading onto shared eigenvector (eigenvectors associated with the 2nd highest eigenvalue for **L** and the with the highest eigenvalue for both **M** matrices).

**Supplementary Figure 7:**
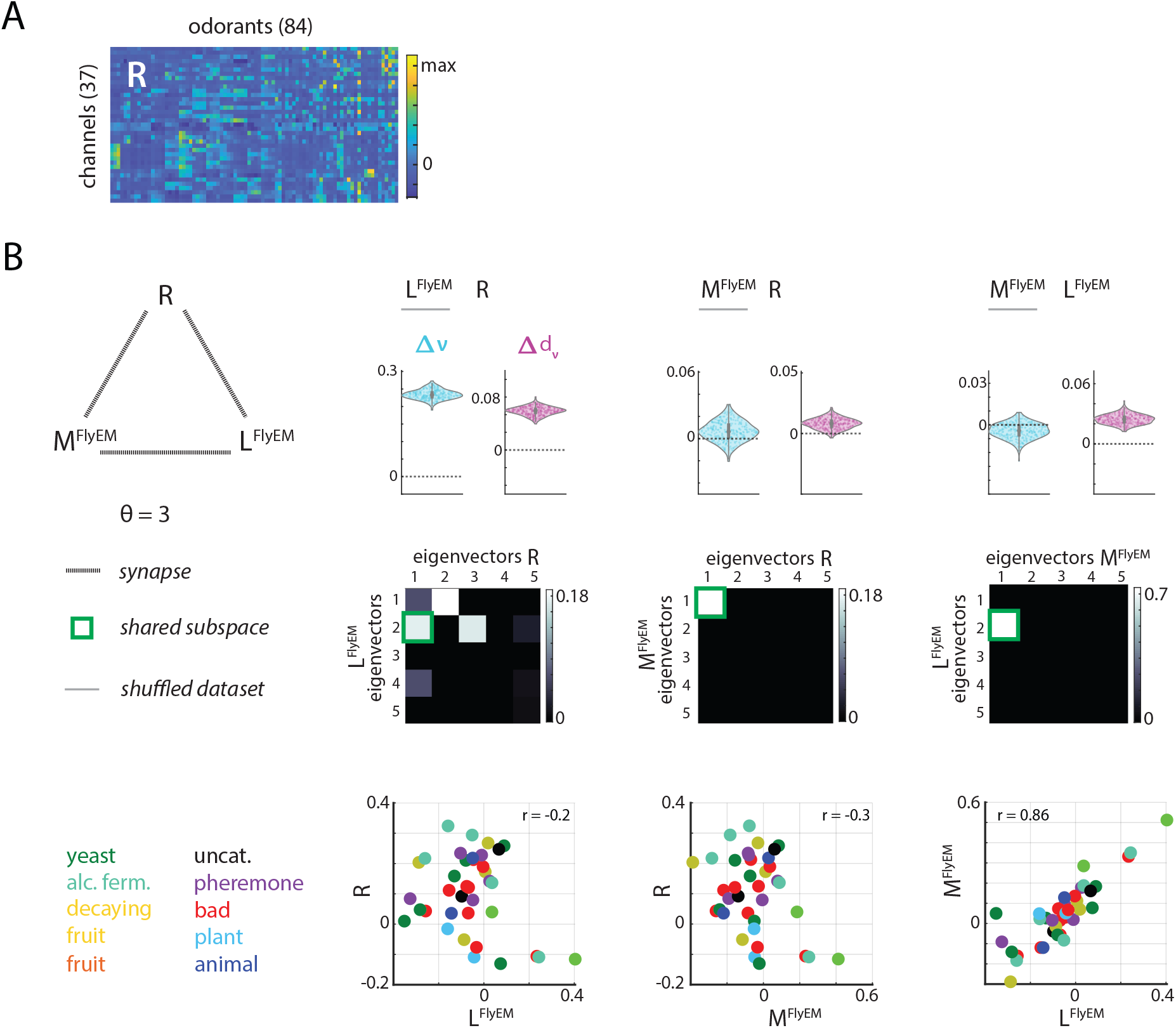
**A**. The response of a subset of 37 olfactory channels to 84 odorants from four categories: monomolecular species, natural odorant blends, concentration series and binary mixtures (matrix **R**) [2]. **B**. Multimodal data analysis with the ENSD. Comparing synaptic count data from *FlyEM* (using the corresponding 37 channels) with odorant response data in **R**. *Top row* : changes in ENSD and distance metric when one dataset is shuffled (underlined in grey). *Second row* : **W** matrices containing eigenvector overlaps corresponding to the top 5 eigenvalues shown, all statistically insignificant overlaps (*p* > 0.05), are set to zero. *Bottom row* : Comparison of loadings onto shared dimension.

